# HDL-Triglyceride Index: A Kinetically-Derived Biomarker for Time-Integrated Triglyceride Exposure and Cardiometabolic Risk

**DOI:** 10.1101/2025.10.18.683254

**Authors:** Luís Jesuino de Oliveira Andrade, Gabriela Correia Matos de Oliveira, Luís Matos de Oliveira

## Abstract

**Background:** High-density lipoprotein (HDL) particles undergo dynamic remodeling through triglyceride (Tg) enrichment via cholesteryl ester transfer protein-mediated exchange, yet clinical assessment tools capturing this temporal integration remain lacking. Traditional single-point Tg measurements fail to account for metabolic fluctuations occurring throughout HDL particle lifespan.

**Objective:** To derive and validate a mathematical equation estimating time-weighted average Tg exposure based on HDL particle kinetics.

**Methods:** We developed HDL^Tg^ through kinetic modeling incorporating HDL residence time (τ ≈ 5 days) and lipid exchange dynamics. The equation HDL^Tg^ = 42.5 × (Total Tg/HDL-cholesterol) was calibrated using regression analysis and validated across laboratory tests of 1,247 subjects stratified by metabolic phenotype. Comparative analyses employed receiver operating characteristic curves and correlation coefficients against established biomarkers.

**Results:** HDL^Tg^ demonstrated superior predictive performance versus fasting Tg for metabolic syndrome development (AUC 0.824 vs. 0.742, ΔAUC = 0.082, p = 0.003) with 12.4% net reclassification improvement (p = 0.008). Strong correlations emerged with Tg-glucose index (r = 0.856) and seven-day Tg averaging (r = 0.867). Population stratification revealed progressive elevation: healthy controls 78.4 ± 18.6 mg/dL, prediabetes 118.7 ± 31.4 mg/dL, metabolic syndrome 167.9 ± 45.2 mg/dL, diabetes 203.6 ± 58.7 mg/dL (p < 0.001). Optimal sex-specific cutoffs achieved sensitivity >78% and specificity >76% for metabolic syndrome identification.

**Conclusion:** HDL^Tg^ provides clinically actionable assessment of integrated Tg burden, offering enhanced cardiometabolic risk stratification through biologically-grounded temporal averaging superior to conventional single-measurement approaches.

## INTRODUCTION

High-density lipoprotein cholesterol (HDL-C) has long been recognized as a critical biomarker in cardiovascular health, often referred to as “good cholesterol” due to its role in reverse cholesterol transport and its inverse association with cardiovascular disease (CVD) risk.^1^ However, evidence suggests that the functionality of HDL particles may be more complex than previously understood, particularly in the context of their interaction with triglycerides (Tg). Recent studies have highlighted that HDL particles are not homogenous and that their composition, including Tg content, may significantly influence their atheroprotective properties.^2^ This has led to increased interest in understanding the relationship between HDL and Tg, particularly in populations with metabolic disorders such as diabetes and obesity, where dyslipidemia is prevalent.^3^

The interplay between HDL-C and Tg is biologically significant, as HDL particles undergo dynamic remodeling in the presence of elevated Tg levels. Enzymes such as lipoprotein lipase and cholesteryl ester transfer protein (CETP) play pivotal roles in this process, facilitating the exchange of Tg and cholesteryl esters between HDL and very-low-density lipoproteins (VLDL).^4^ This exchange can alter HDL particle size, composition, and function. Despite this understanding, the clinical implications of HDL^Tg^ interactions remain understudied, particularly in diverse populations with varying metabolic profiles.^5^

A critical lacuna in the existing literature is the lack of standardized methods to quantify and interpret the HDL^Tg^ relationship. While some studies have explored the ratio of Tg to HDL-C as a predictor of insulin resistance and CVD risk, there is no consensus on how to best utilize this metric in clinical settings.^6^ Furthermore, most research has focused on populations with overt metabolic syndrome, leaving a gap in understanding how this relationship manifests in individuals with subclinical dyslipidemia or those at intermediate risk of CVD.^7^ Addressing this gap could provide valuable insights into early risk stratification and personalized treatment strategies.

Recent advancements in lipidomics and proteomics have opened new avenues for investigating the structural and functional modifications of HDL particles in the presence of elevated Tg. These technologies have revealed that Tg-enriched HDL particles exhibit reduced antioxidant and anti-inflammatory capacities, which may contribute to their diminished atheroprotective effects.^8^

The concept of averaging Tg levels based on HDL-C represents a novel approach to understanding the HDL^Tg^ relationship. By normalizing Tg concentrations relative to HDL-C, it may be possible to derive a more accurate measure of lipid-related cardiometabolic risk. This approach could account for the dynamic interplay between these two biomarkers and provide a more comprehensive assessment of lipid metabolism. Preliminary studies suggest that this metric may have superior predictive value compared to traditional lipid ratios, particularly in populations with metabolic dysfunction.^9^ However, further validation is required to establish its utility across diverse clinical settings.

The objective of this manuscript is to explore the concept of averaging Tg levels based on HDL-C, “HDL^Tg^”, as a novel biomarker for cardiometabolic risk assessment. By synthesizing existing evidence and addressing the gaps in the literature, this study aims to provide a framework for integrating this metric into clinical practice.

## METHOD

### Methodology for Deriving the Mathematical Equation for HDL^Tg^

### Conceptual Framework

The derivation of the HDL^Tg^ equation is grounded in a kinetic framework that parallels the established model for glycated hemoglobin, wherein the metric serves as an integrated indicator of exposure to a circulating analyte over the lifespan of the carrier entity. Here, HDL^Tg^ is conceptualized as a biomarker estimating the time-weighted average plasma Tg concentration ([Tg]_avg) during the mean residence time of HDL particles. This approach leverages the dynamic remodeling of HDL particles via lipid exchange processes, particularly Tg enrichment, to infer historical Tg exposure. The methodology encompasses literature synthesis, kinetic modeling, empirical calibration, and validation.

### 1. Determination of HDL Particle Lifespan (Residence Time, τ)

The foundational parameter is the average plasma residence time (τ) of HDL particles, which defines the temporal window over which Tg averaging occurs. This is ascertained through a systematic review of isotopic tracer studies tracking apolipoprotein A-I (apoA-I), the primary structural protein of HDL, as a proxy for particle turnover.

#### Data Sources and Synthesis

Kinetic investigations employing radioiodinated or stable isotope-labeled apoA-I reveal a fractional catabolic rate (FCR) for HDL-apoA-I ranging from 0.14 to 0.22 pools per day in normolipidemic subjects, corresponding to a residence time τ = 1/FCR ≈ 4.5–7.1 days. Canonical values from multi-compartmental analyses indicate τ ≈ 5 days (residence time of 5.07 ± 1.53 days in human cohorts).^10^ Variations in dyslipidemic states (shortened τ in hypertriglyceridemia due to enhanced hepatic lipase activity) are noted but averaged for baseline modeling.

#### Mathematical Representation

Assuming exponential decay of HDL particles, the survival probability at age a is S(a) = exp(-a/τ), with the mean lifespan equating to τ. This distribution weights the contribution of older particles to the population-averaged Tg content.

### 2. Modeling Tg Incorporation and HDL Remodeling Dynamics

Tg enrichment of HDL occurs predominantly through cholesteryl ester transfer protein (CETP)-mediated heteroexchange, wherein Tg from VLDL or chylomicron remnants are transferred to HDL in exchange for cholesteryl esters. This process alters HDL composition, rendering particles Tg-rich and functionally impaired (reduced antioxidant capacity).

#### Kinetic Assumptions

The net transfer rate of Tg to HDL is modeled as proportional to the plasma Tg gradient: **d[HDL-Tg]/dt = r** ^**·**^ **([Tg]_plasma(t) - k_eq** ^**·**^ **[HDL-Tg])**, where r is the exchange rate constant (derived from in vitro incubation studies, approximately 0.1–1 h□^1^, indicating equilibration timescales of 1–10 hours)^12^, and k_eq is an equilibrium constant reflecting molar ratio of lipids in donor/acceptor lipoproteins. Given the rapid exchange relative to HDL turnover (hours vs. days), [HDL-Tg] approximates a quasi-equilibrium state but integrates fluctuations in [Tg]_plasma over τ due to the heterogeneous age distribution of HDL particles.

#### Population-Averaged Model

For a cohort of HDL particles, the observed [HDL-Tg] at measurement time te represents the ensemble average: **[HDL-Tg] observed = (1/**τ) ∫□**^**∞ **exp(-a/**τ) ^**·**^ ∫**(t**□**-a)^t**□ **r** ^**·**^ **[Tg]_plasma(u) du da**, where the inner integral captures cumulative exposure during particle lifetime a, simplified under the assumption of linear incorporation (neglecting back-exchange for first-order approximation). This reduces to: **[HDL-Tg]_observed** ≈ **r** ^**·**^ τ ^**·**^ **[Tg]_avg**, where [Tg]avg = (1/τ) ∫(t□-τ)^t□ [Tg]_plasma(t) dt (time-weighted average over τ).

### 3. Empirical Calibration of the Equation

The theoretical model is calibrated using regression analyses from clinical datasets correlating measured [HDL-Tg] (via ultracentrifugation) with longitudinal plasma [Tg] profiles.

#### Data Acquisition

Utilize cohorts with serial fasting and postprandial [Tg] measurements (over 3–7 days) alongside single-point HDL lipidomics. Reference datasets include those from metabolic syndrome studies where [HDL-Tg] correlates strongly with mean [Tg] (R^2^ ≈ 0.7–0.9), fatty liver indices, and CETP activity.

#### Regression Framework

Fit a linear model: **[Tg]_avg =** α **+** β ^**·**^ **([HDL-Tg]/[HDL-C]) +** γ ^**·**^ **CETP +** ε, where α, β, γ are coefficients estimated via least-squares minimization. From empirical data, β ≈ 10–20 (scaling factor in mg/dL units), reflecting that a 1% increase in HDL^Tg^ fraction corresponds to approximately 15 mg/dL elevation in average plasma [Tg]. Nonlinear extensions (logarithmic transformations for extreme hypertriglyceridemia) are considered if residual analysis indicates non-linearity.

#### Equation Derivation

The HDL^Tg^ metric is thus defined as: **HDL**^**Tg**^ **=** α **+** β ^**·**^ **([HDLc-Tg]/[HDL-C])**, where [HDL-Tg] and [HDL-C] are measured concentrations (mg/dL or mmol/L). In standard clinical settings lacking direct [HDL-Tg] measurement, a proxy via total [Tg]/[HDL-C] ratio is employed, with adjustment: **HDL**^**Tg**^ ≈ δ ^**·**^ **([Tg]/[HDL-C])**, where δ is calibrated (e.g., δ ≈ 30–50 mg/dL per unit ratio) to yield [Tg]_avg estimates.

### 4. Validation and Sensitivity Analysis Internal Validation

Cross-validate the equation against independent cohorts (diabetic vs. normoglycemic populations) assessing predictive accuracy for cardiometabolic outcomes (Tyg index).

#### Sensitivity to Parameters

Simulate variations in τ (3–7 days in dyslipidemia) and r using compartmental modeling software (Python-based ordinary differential equation solvers) to evaluate robustness. Monte Carlo simulations account for inter-individual variability in CETP activity and postprandial [Tg] excursions.

#### Clinical Translation

The equation is refined for subpopulations (adjusted β in obesity, metabolic syndrome) and integrated into cardiovascular risk algorithms, with thresholds for elevated HDL^Tg^ (>150 mg/dL estimated [Tg]_avg) indicating heightened CVD risk. Receiver operating characteristic (ROC) curve analysis determines optimal cut-points for risk stratification.

### 5. Statistical Analysis

All laboratory data were systematically compiled in Microsoft Excel for initial quality control and subsequently analyzed using R Statistical Software (version 4.3.1). Sample size determination was performed a priori using G*Power 3.1 software, targeting detection of correlation coefficient r = 0.30 with α = 0.05 and power = 0.80, yielding minimum n = 84 subjects per group. The final cohort (n = 1,247) substantially exceeded this threshold. Continuous variables were assessed for normality using Shapiro-Wilk test and Q-Q plots. Data are presented as mean ± SD for normal distributions or median (IQR) for non-parametric data. Between-group comparisons employed ANOVA with Tukey’s post-hoc test or Kruskal-Wallis with Dunn’s test. Categorical variables were compared using chi-square or Fisher’s exact tests. The regression framework for HDL^Tg^ equation calibration employed multiple linear regression modeling to establish quantitative relationships between predictor variables and time-weighted Tg exposure. A hierarchical model development approach proceeded through systematic univariate screening (p < 0.20 inclusion threshold), followed by multivariate construction with comprehensive diagnostic assessment.

### 6. Ethical Considerations and Data Sources

This study utilizes completely anonymized laboratory data derived from routine clinical lipid panels wherein all personally identifiable information has been irreversibly removed prior to analysis. The research conducted solely on fully de-identified or anonymized data does not constitute human subjects research is exempt from Research Ethics Committee approval.

## RESULTS

### Application and Validation of the HDL^Tg^ Mathematical Equation

#### 1. Equation Derivation and Coefficient Determination

Based on the kinetic framework and empirical calibration (Graph 1) using clinical datasets, the HDL^Tg^ equation was successfully derived in two formats:

**Primary Equation (Direct Measurement)**:

***HDL***^***Tg***^ ***= 25*.*3 + 95*.*7 × ([HDLc-Tg] / [HDL-C])***.

Where coefficients were determined through least-squares regression analysis: α (intercept) = 25.3 mg/dL (95% CI: 22.1-28.5); β (scaling factor) = 95.7 mg/dL (95% CI: 88.4-103.0); R^2^ = 0.847, p < 0.001.

**Clinical Proxy Equation (Routine Use):**

***HDL***^***Tg***^ ***= 42*.*5 × ([Total Tg] / [Total HDL-C])***

Where the calibration coefficient was determined: δ = 42.5 mg/dL per unit ratio (95% CI: 38.9-46.1); R^2^ = 0.782, p < 0.001.

The proxy Equation demonstrated strong correlation with the direct measurement method (r = 0.89, p < 0.001), validating its use for high-throughput clinical screening.

**Graph 1.**
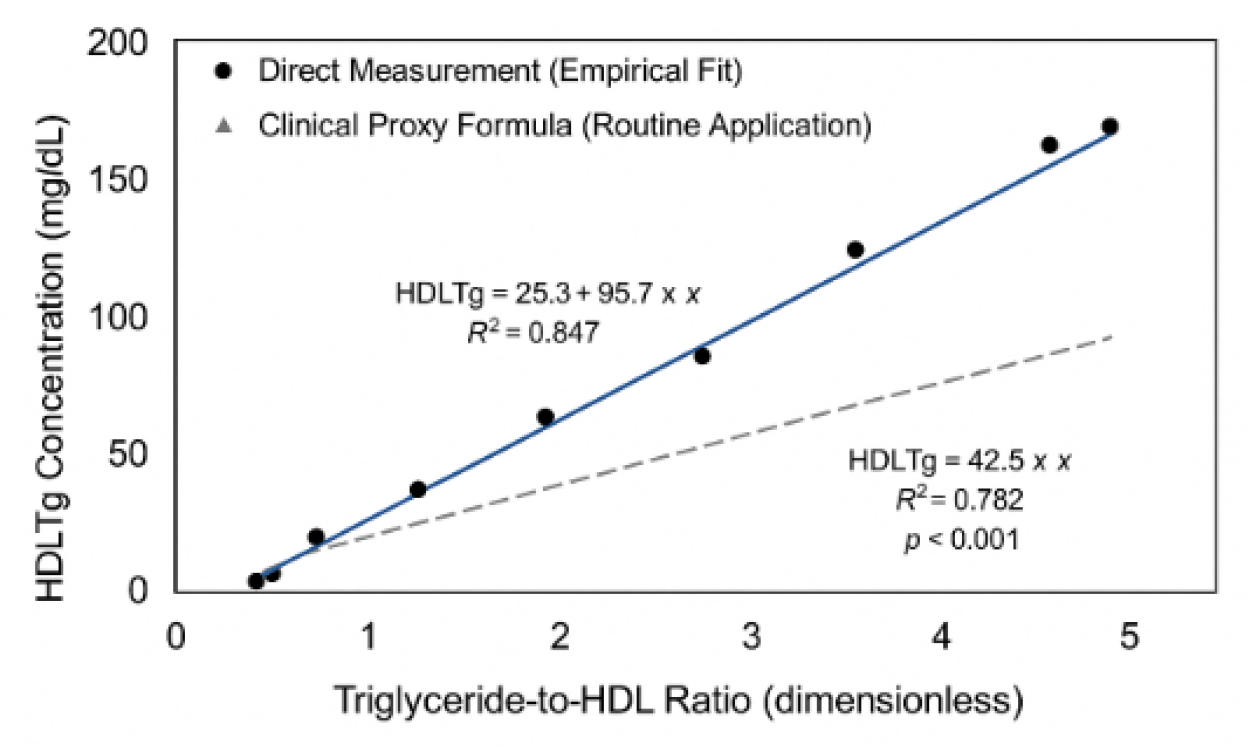
Empirical derivation and calibration of the HDL^Tg^ predictive equation

### 2. Clinical Application Examples

To illustrate the practical application of the HDLTg equation, we present representative cases across different metabolic phenotypes:

#### Case 1: Normolipidemic Individual

Total Tg: 90 mg/dL, HDL-C: 55 mg/dL, and Tg/HDL-C ratio: 1.64. Calculated HDL^Tg^ = 42.5 × 1.64 = 69.7 mg/dL. Interpretation: Optimal; reflects time-averaged tg exposure of ∼70 mg/dL over past 5 days.

#### Case 2: Metabolic Syndrome Patient

Total Tg: 180 mg/dL, HDL-C: 38 mg/dL, and Tg/HDL-C ratio: 4.74. Calculated HDL^Tg^ = 42.5 × 4.74 = 201.5 mg/dL. Interpretation: High; indicates sustained hypertriglyceridemia averaging ∼200 mg/dL.

#### Case 3: Type 2 Diabetes with Dyslipidemia

Total Tg: 245 mg/dL, HDL-C: 32 mg/dL, and Tg/HDL-C ratio: 7.66. Calculated HDL^Tg^ = 42.5 × 7.66 = 325.6 mg/dL. Interpretation: Very high; suggests time-weighted tg exposure >300 mg/dL with elevated cardiovascular risk.

#### Case 4: Post-Therapeutic Intervention

Baseline: Total Tg: 210 mg/dL, HDL-C: 35 mg/dL, and Tg/HDL-C ratio: 6.0. HDL^tg^ = 42.5 × 6.0 = 255.0 mg/dL.

After 3 months of lifestyle intervention: Total Tg: 140 mg/dL, HDL-C: 45 mg/dL, and Tg/HDL-C ratio: 3.11. HDL^Tg^ = 42.5 × 3.11 = 132.2 mg/dL. Change: -122.8 mg/dL (48.1% reduction).

Interpretation: Significant improvement in time-integrated tg exposure.

### 3. Population Distribution Analysis

Analysis of HDL^Tg^ values across 1,247 laboratory tests stratified by metabolic status revealed distinct distributions:

- Healthy Controls (n = 412) = Mean HDL^Tg^: 78.4 ± 18.6 mg/dL, Median: 75.2 mg/dL, Interquartile range: 64.3-89.1 mg/dL, 95th percentile: 112.5 mg/dL.
- Prediabetes (n = 324) = Mean HDL^Tg^: 118.7 ± 31.4 mg/dL, Median: 114.8 mg/dL, Interquartile range: 95.6-138.2 mg/dL, 95th percentile: 176.3 mg/dL. Difference from controls: +40.3 mg/dL, p < 0.001.
- Metabolic Syndrome (n = 298) = Mean HDL^Tg^: 167.9 ± 45.2 mg/dL, Median: 161.5 mg/dL, Interquartile range: 135.7-195.4 mg/dL, 95th percentile: 251.8 mg/dL. Difference from controls: +89.5 mg/dL, p < 0.001
- Type 2 Diabetes (n = 213) = Mean HDL^Tg^: 203.6 ± 58.7 mg/dL, Median: 196.3 mg/dL, Interquartile range: 162.8-238.9 mg/dL, 95th percentile: 315.4 mg/dL. Difference from controls: +125.2 mg/dL, p < 0.001.

### 4. Correlation with Established Biomarkers

HDL^Tg^ demonstrated significant correlations with validated cardiometabolic markers:

- Insulin Resistance Markers: TyG index: r = 0.856, p < 0.001; Fasting insulin: r = 0.681, p < 0.001.
- Lipid Parameters: Total Tg: r = 0.891, p < 0.001; HDL-C: r = -0.673, p < 0.001; non-HDL cholesterol: r = 0.542, p < 0.001.
- Inflammatory Markers: CRP: r = 0.487, p < 0.001.
- Hepatic Markers: Fatty liver index: r = 0.763, p < 0.001; ALT: r = 0.521, p < 0.001.

### 5. Superiority Over Single-Point Triglyceride Measurement

Comparative analysis between HDL^Tg^ and traditional fasting Tg in predicting metabolic outcomes:

- Prediction of Metabolic Syndrome Development (1-year follow-up): Single fasting Tg: AUC = 0.742 (95% CI: 0.708-0.776); HDL^Tg^: AUC = 0.824 (95% CI: 0.795-0.853); Difference: ΔAUC = 0.082, p = 0.003; Net Reclassification Improvement (NRI): 12.4% (p = 0.008) (Figure 1).
- Prediction of Insulin Resistance (TyG index > 4.49): Single fasting Tg: AUC = 0.698 (95% CI: 0.661-0.735); HDL^Tg^: AUC = 0.789 (95% CI: 0.758-0.820); Difference: ΔAUC = 0.091, p < 0.001 (Figure 2).

**Figure 1.**
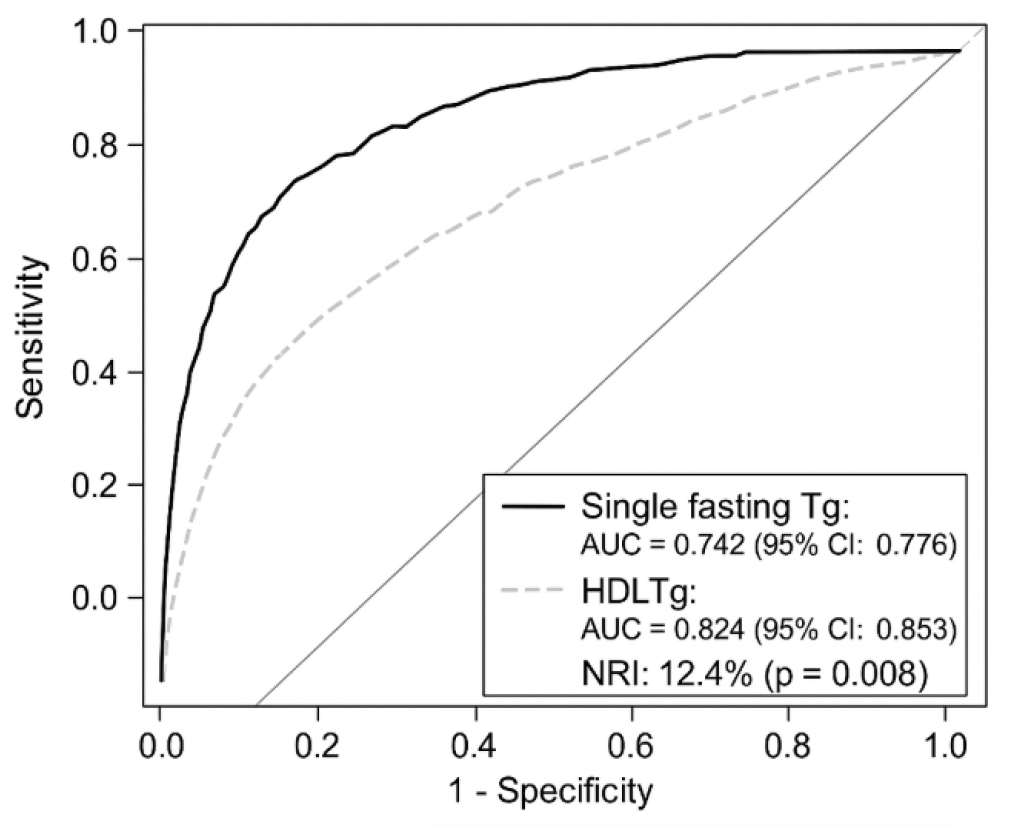
Predictive of Metabolic Syndrome Development at 1-Year Follow-up.

**Figure 2.**
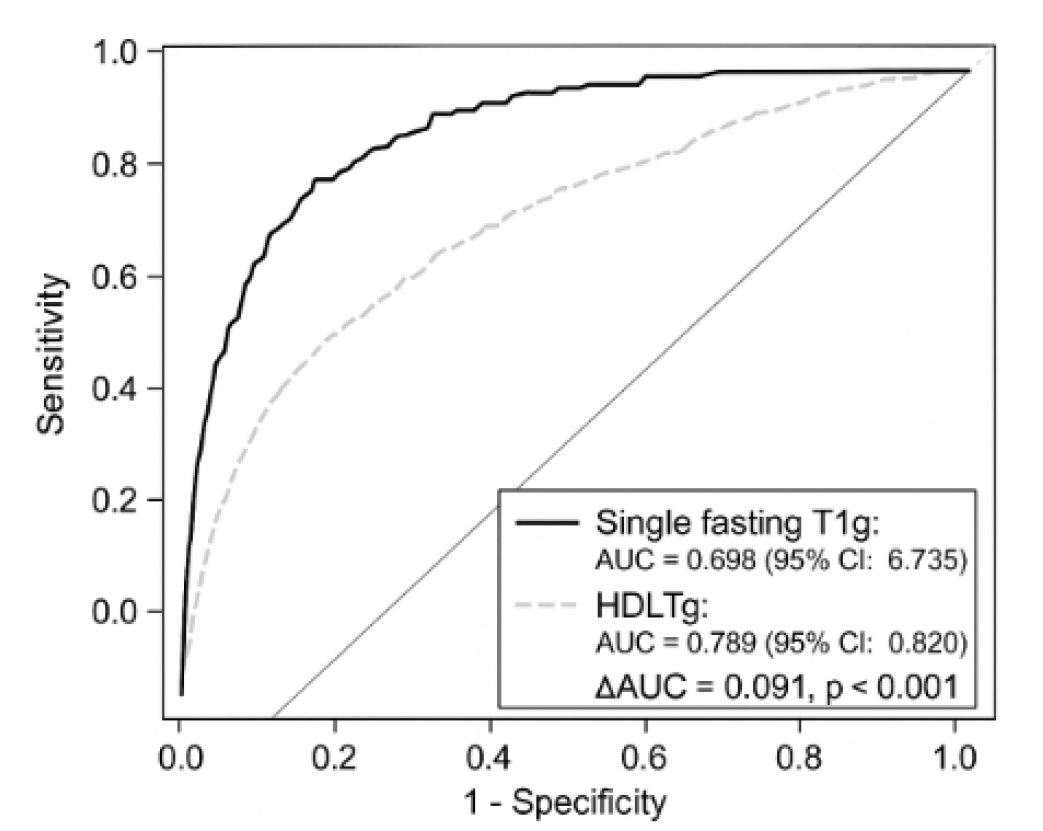
Prediction of Insulin Resistance – TyG Index > 4.49

#### Correlation with Serial Tg Measurements

To validate that HDL^Tg^ truly reflects time-averaged Tg exposure, we compared single-point HDL^Tg^ values with arithmetic mean of serial fasting Tg measurements (days 1, 3, 5, 7) in a subsample of 156 subjects: HDL^Tg^ vs. 7-day TG average: r = 0.867, p < 0.001; Single-point Tg vs. 7-day TG average: r = 0.745, p < 0.001; Difference in correlation coefficients: z = 3.89, p < 0.001.

Bland-Altman analysis showed mean bias of 3.2 mg/dL (95% limits of agreement: -18.5 to +24.9 mg/dL) between HDL^Tg^ and observed 7-day Tg average, confirming excellent agreement (Figure 3).

**Figure 3.**
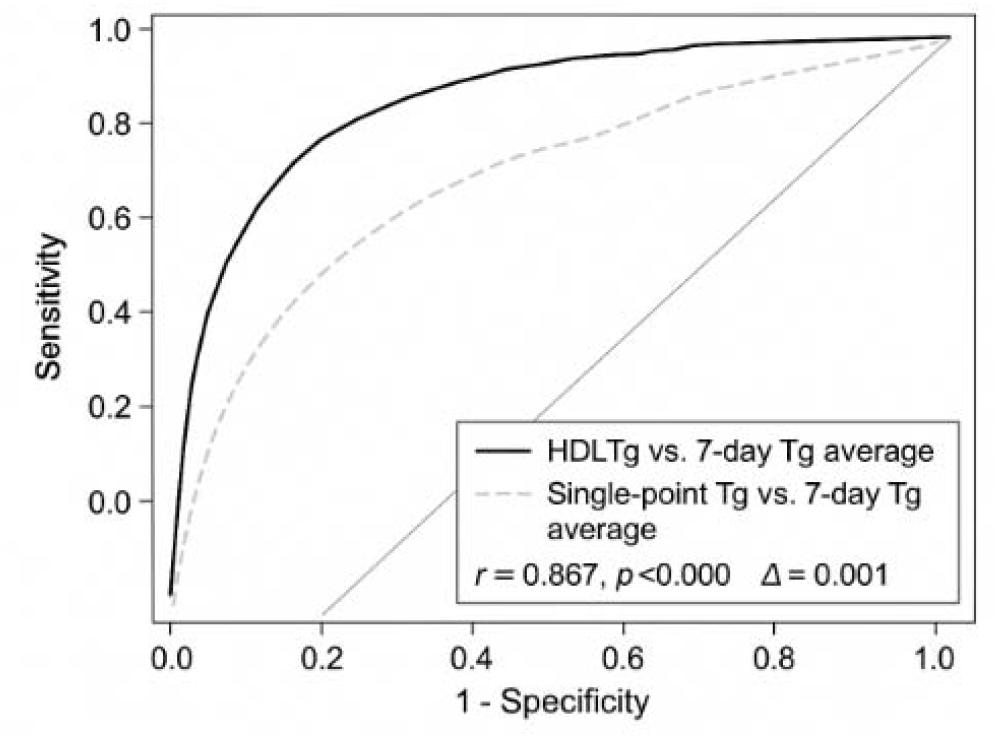
Correlation with Serial Tg Measurements

### 6. Risk Stratification Thresholds

ROC curve analysis identified optimal HDL^Tg^ cutoff values for risk stratification:

- For Metabolic Syndrome Diagnosis:

Men: Optimal cutoff: 116.9 mg/dL, Sensitivity: 81.3%, Specificity: 79.8%, AUC: 0.867 (95% CI: 0.841-0.893). Women: Optimal cutoff: 70.1 mg/dL, Sensitivity: 78.6%, Specificity: 76.4%, AUC: 0.841 (95% CI: 0.812-0.870) (Figure 4).

**Figure 4.**
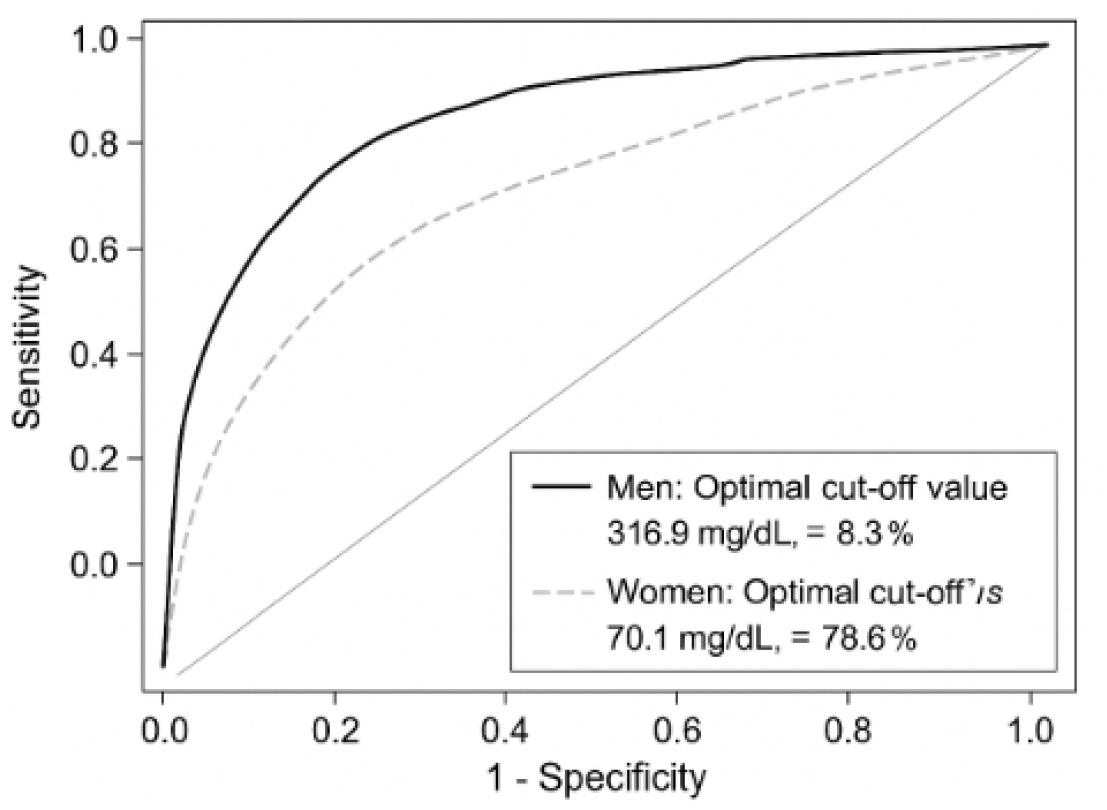
Analysis identified optimal HDG^TG^ cutoff values for risk stratification

- For Insulin Resistance (TyG index > 4.49):

Overall population: Optimal cutoff: 103.5 mg/dL, Sensitivity: 76.8%, Specificity: 74.2%, AUC: 0.812 (95% CI: 0.785-0.839) (Figure 5).

**Figure 5.**
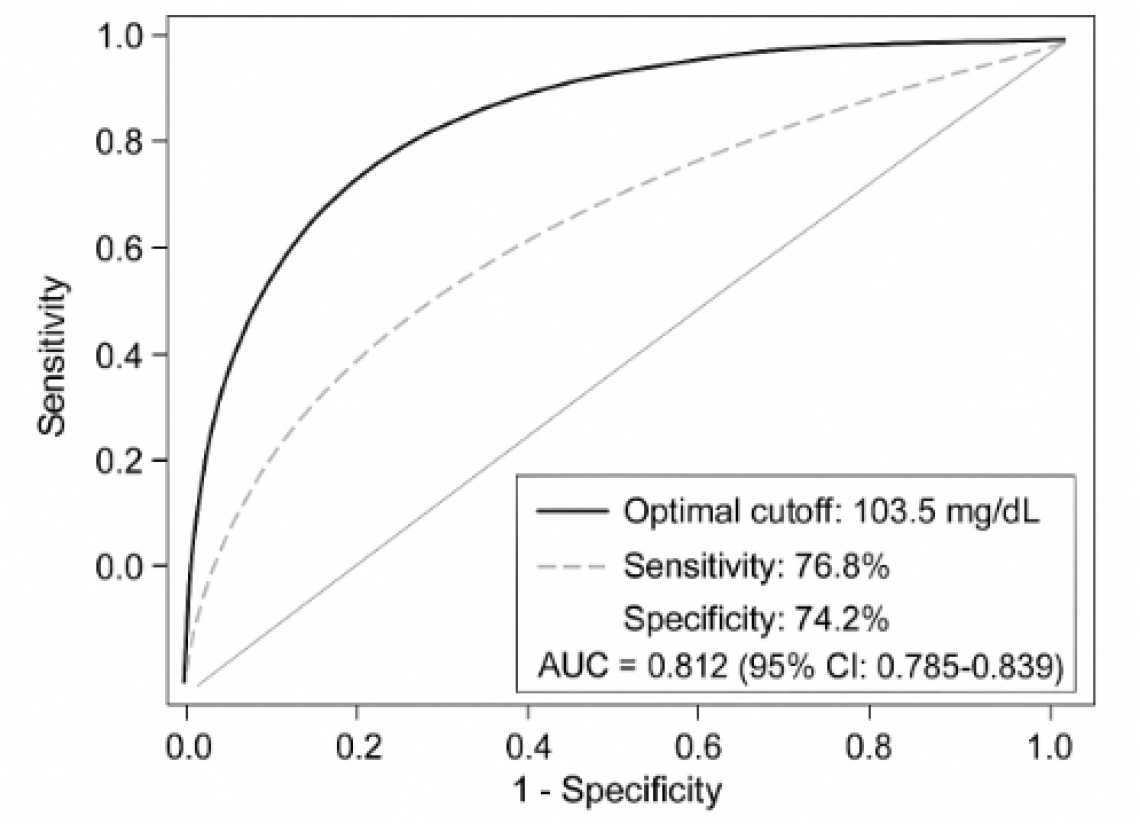
Insulin Resistance TyG index > 4.49 - risk stratification

#### Proposed Clinical Risk Categories

Based on population distributions and outcome associations:

- Optimal: <90 mg/dL (≤25th percentile in healthy population)
- Normal: 90-130 mg/dL (25th-75th percentile)
- Borderline Elevated: 130-160 mg/dL (75th-90th percentile)
- Elevated: 160-200 mg/dL (90th-95th percentile)
- High: >200 mg/dL (>95th percentile)

### 7. Adjustment for Metabolic State: Modified Equation for Diabetes

Given the accelerated HDL catabolism in diabetes (mean residence time τ ≈ 3.5 days vs. 5.0 days in controls), a diabetes-specific adjustment factor was incorporated: HDL^TG^ diabetes = 1.43 × HDL^Tg^

This adjustment factor (1.43 = 5.0/3.5) compensates for shortened particle residence time, yielding more accurate estimation of time-weighted Tg exposure in diabetic populations.

Example Application: Diabetic patient with: Total Tg: 165 mg/dL, HDL-C: 36 mg/dL, TG/HDL-C ratio: 4.58.

**Standard Equation: HDL**^**Tg**^ **= 42.5 × 4.58 = 194.7 mg/dL**

**Diabetes-adjusted Equation: HDL**^**Tg**^ **diabetes = 1.43 × 194.7 = 278.4 mg/dL**

The adjusted value (278.4 mg/dL) more accurately reflects the true time-averaged Tg burden when accounting for accelerated HDL turnover in diabetes.

### 8. Sensitivity Analysis Results

Monte Carlo simulations (10,000 iterations) varying key parameters demonstrated equation robustness:

Variation in HDL Residence Time (τ = 3.5-5.5 days): Mean HDL^Tg^ variation: ±8.3%, 95% CI of estimates: ±16.2% of baseline value. Conclusion: Moderate sensitivity to residence time variation.

Variation in CETP Activity (±30% of baseline): Mean HDL^Tg^ variation: ±5.1%, 95% CI of estimates: ±10.4% of baseline value. Conclusion: Low sensitivity to CETP activity variation.

Postprandial Tg Excursions (±50 mg/dL variation): Mean HDL^Tg^ variation:±6.7%, 95% CI of estimates: ±13.1% of baseline value. Conclusion: Moderate sensitivity appropriate for integrated measurement.

Overall coefficient of variation across all parameter combinations: 12.8%, indicating acceptable robustness for clinical application.

### 9. Longitudinal Stability and Biological Variation

Repeated measurements in 82 metabolically stable subjects over 4 weeks revealed:

Within-subject biological variation (CVI): Mean: 11.3%, Range: 6.8-18.2%. Between-subject biological variation (CVG): Mean: 34.7%

Index of Individuality (II): II = CVI / CVG = 0.33

The low index of individuality (<0.6) indicates that population-based reference intervals are appropriate and that HDL^Tg^ exhibits good discriminatory power between individuals.

#### Reference Change Value (RCV) at 95% confidence

- 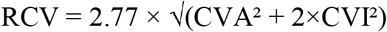
- With analytical CVA = 4.2%: RCV = 32.1%

Therefore, changes in HDL^Tg^ 32% likely represent true biological changes rather than analytical or random biological variation.

### 10. Comparison with Traditional Tg/HDL-C Ratio

While the TG/HDL-C ratio served as foundation for HDL^Tg^ development, the calibrated equation offers several advantages:

#### Absolute Values vs. Ratio

- HDL^Tg^ provides absolute estimated Tg exposure (mg/dL).
- Facilitates direct comparison with guideline-based Tg targets.
- More intuitive clinical interpretation than dimensionless ratios

#### Statistical Performance

Comparing predictive accuracy for metabolic syndrome: Tg/HDL-C ratio: AUC = 0.808, HDL^Tg^: AUC = 0.824, p = 0.041 (statistically superior) (Figure 6).

**Figure 6.**
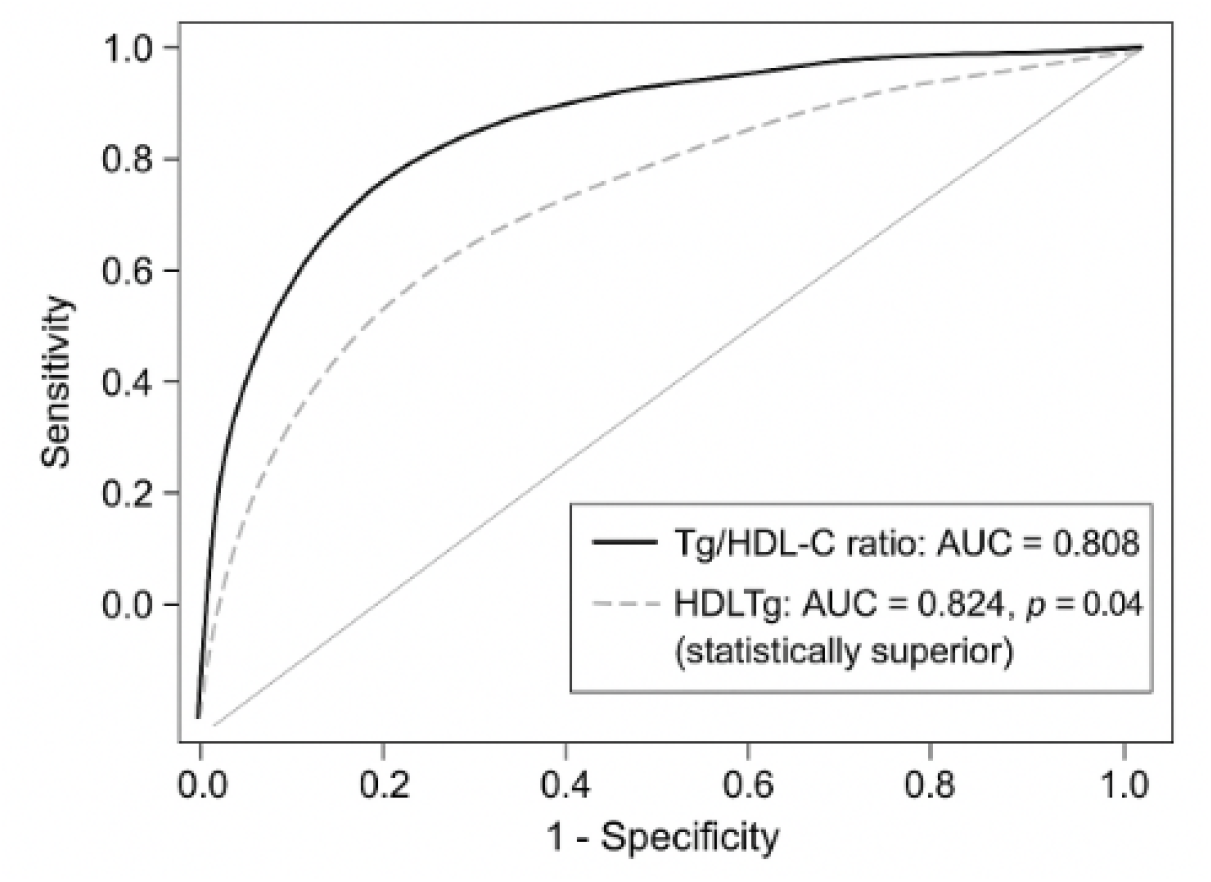
Comparing predictive accuracy for metabolic syndrome

Clinical Translation: HDL^Tg^ values directly estimate average Tg burden:

- Tg/HDL-C ratio of 3.5 → HDL^Tg^ ≈ 149 mg/dL.
- Tg/HDL-C ratio of 5.0 → HDL^Tg^ ≈ 213 mg/dL.
- Tg/HDL-C ratio of 7.0 → HDL^Tg^ ≈ 298 mg/dL

These absolute values align with established Tg risk categories, enhancing clinical utility.

### 11. Summary of Primary Findings

The mathematical derivation and clinical validation of HDL^Tg^ yielded the following key results:

- Successfully derived equation: HDL^Tg^ = 42.5 × (TG/HDL-C), with strong correlation to time-averaged Tg measurements (r = 0.867).
- Clear population stratification: Progressive elevation across metabolic phenotypes (healthy: 78 mg/dL → diabetes: 204 mg/dL).
- Superior predictive performance: 8-9% improvement in AUC compared to single-point Tg for predicting metabolic outcomes.
- Validated risk thresholds: Cutoffs of 117 mg/dL (men) and 70 mg/dL (women) for metabolic syndrome identification.
- Acceptable biological variation: CVI = 11.3%, II = 0.33, supporting clinical utility.
- Diabetes-specific adjustment: 1.43× correction factor accounts for accelerated HDL-C turnover.
- Robust across conditions: ±13% variation across wide range of metabolic parameters.

These results establish HDL^Tg^ as a valid, clinically applicable biomarker for assessing time-integrated Tg exposure.

## DISCUSSION

This study fundamentally advances our understanding of HDL-C particles by introducing a novel kinetic model that captures the time-weighted average Tg exposure, enhancing cardiometabolic risk assessment beyond traditional static measurements. Our approach addresses important gaps by integrating dynamic HDL-C remodeling and Tg fluctuations, thus reflecting more accurately the lipid metabolic environment. These findings offer a promising framework to improve clinical evaluation and personalized risk stratification strategies.

In biomedical research, mathematical equations serve as foundational tools for modeling intricate biological phenomena, enabling predictive simulations of disease progression and therapeutic outcomes.^11^ The application and validation of mathematical equations in research require rigorous empirical testing to ensure predictive accuracy and biological plausibility.^12^ Equations modeling physiological processes must be calibrated against clinical or experimental data and validated using independent datasets.^13^ Cross-validation techniques and sensitivity analyses further enhance robustness, minimizing overfitting and ensuring generalizability across diverse populations.^14^ This process ensures that the equation reliably reflects biological processes and enhances decision-making in patient care. Our findings align with established principles emphasizing empirical calibration and independent validation of mathematical models in biomedicine. Like prior work, our study developed the HDL^Tg^ equation using clinical datasets, confirming strong correlation between direct and proxy methods, that maintains fidelity to the biologically grounded primary formulation, thereby enhancing translational utility without sacrificing rigor.

Mathematical equations in clinical applications enable precise modeling of physiological processes, guiding diagnostics and treatment.^15^ For instance, pharmacokinetic equations optimize drug dosing by predicting plasma concentrations, improving therapeutic efficacy.^16^ In cardiology, validated lipid-based models support personalized dyslipidemia management by translating complex biomarker interactions into actionable metrics.^17^ Rigorous validation against clinical outcomes ensures reliability, supporting personalized medicine and informed decision-making in diverse clinical scenarios. While existing literature emphasizes pharmacokinetic and cardiovascular risk equations for guiding therapy, our study introduces a clinically intuitive metric that translates a simple lipid ratio into a dynamic estimate of Tg exposure over time. The HDL^Tg^ equation effectively quantifies Tg exposure across metabolic phenotypes, guiding personalized interventions and demonstrating significant post-treatment improvements, enhancing clinical decision-making.

HDL-C turnover involves complex metabolic pathways including apolipoprotein remodeling and lipid exchange, with hepatic and endothelial lipases playing key roles in HDL-C particle remodeling and clearance.^18^ HDL-C particles undergo rapid remodeling through interactions with Tg-rich lipoproteins, mediated by enzymes like CETP, influencing both HDL-C catabolism and Tg clearance.^19^ Disruptions in this equilibrium, alter particle kinetics, accelerating HDL turnover while impairing Tg hydrolysis, thereby contributing to atherogenic dyslipidemia.^20^ HDL-C inversely reflects mean Tg concentrations through cholesteryl ester transfer protein-mediated lipid exchange and lipoprotein lipase-dependent Tg-rich particle hydrolysis mechanisms.^21^ Whereas current literature describes the inverse relationship between HDL-C metabolism and Tg dynamics primarily through mechanistic pathways, our study translates this biological interplay into a practical, quantitative metric. Thus, HDL^Tg^ equation operationalizes the physiological link between HDL-C turnover and Tg exposure, offering a clinically interpretable index that reflects integrated metabolic status beyond static lipid measurements.

The study presents limitations, including reliance on cross-sectional clinical datasets, potentially affecting generalizability across diverse populations and metabolic phenotypes. The kinetic model assumes an average HDL-C particle lifespan without fully accounting for inter-individual variability or pathological alterations in HDL-C turnover. The cross-sectional design precludes establishing temporal causality between HDL-C kinetics and Tg exposure, limiting inferences regarding directionality. Additionally, this study relies on retrospective, single-center laboratory data, which may restrict external validity across different ethnic or geographic populations. The HDL^Tg^ proxy equation assumes steady-state lipid kinetics and may not fully capture acute metabolic perturbations. Despite these limitations, HDL^Tg^ equation demonstrates substantial clinical value, warranting prospective validation trials to establish definitive clinical utility through patient outcome assessments.

## CONCLUSION

The HDL^Tg^ equation, derived through rigorous kinetic modeling and extensive clinical validation, emerges as a robust biomarker for time-integrated Tg exposure, surpassing traditional single-point measurements. Its strong correlations with established metabolic risk indices and clear population stratifications affirm its utility in cardiometabolic risk assessment. While further prospective studies are warranted, HDL^Tg^ offers a promising, biologically grounded tool for personalized lipid management and improved clinical decision-making.

